# Intraspecific variation of thermal tolerance in freshwater insects along elevational gradients: the case of a widespread diving beetle

**DOI:** 10.1101/2024.03.04.583263

**Authors:** Susana Pallarés, José Antonio Carbonell, Félix Picazo, David T. Bilton, Andrés Millán, Pedro Abellán

## Abstract

Species distributed along wide elevational gradients are likely to experience local adaptation and exhibit high plasticity of thermal tolerance traits, as these gradients are characterised by steep environmental changes over short geographic distances (i.e., strong selection differentials). However, the prevalence of adaptive clinal intraspecific variation in thermal tolerance with elevation remains unclear, and this aspect has been poorly studied in freshwater insects. We explored variation in upper (heat coma temperature) and lower (supercooling point) thermal limits and acclimation capacity among Iberian populations of the widespread aquatic beetle *Agabus bipustulatus* (fam. Dytiscidae) across a 2,000-m elevational gradient, from lowland to alpine areas. As minimum, maximum and mean temperatures decline with elevation, we predicted that higher elevation populations will show lower heat tolerances and higher cold tolerances. We also explored whether acclimation capacity is positively related with climatic variability across different elevations. We found significant variation in upper and lower thermal limits among populations of *A. bipustulatus*, but no evidence of local adaptation to different thermal conditions along the altitudinal gradient, as relationships between thermal limits and elevation or climatic variables were in general not significant. Plasticity of upper and lower thermal limits was overall and consistently low in all populations. These results suggest conservatism of the thermal niche, which might be the result of gene flow counteracting the effects of divergent selection, or adaptations in other traits that buffer the exposure of populations to climate extremes. The limited adaptive potential and plasticity of thermal tolerance found here for *A. bipustulatus* imply that even generalist species, distributed along wide environmental gradients, may have little resilience to global warming.

## INTRODUCTION

Under changing environmental conditions, phenotypic plasticity enables organisms to retain their genotypes while generating different phenotypes (Chevin & Hoffmann, 2017). Also, in cases where the environment imposes substantial constraints and there is some capacity for adaptation within populations, natural selection can promote genetic changes that enhance the fitness of individuals within their specific local environments (Kawecki & Ebert, 2004). For example, thermal traits and their plasticity frequently vary among populations of the same species, typically mirroring changes in the thermal environment (Freidenburg & Skelly, 2004; Sinclair et al., 2012, 2016). The study of population variation and phenotypic plasticity in thermal traits is essential to accurately predict species responses in the face of climate change (Valladares et al., 2014; DeMarche et al., 2019). Indeed, assuming that all populations of a species have identical tolerance ranges and acclimation capacities could overestimate its real ability to cope with environmental changes, especially for widespread species (O’Neill et al., 2008; Naya & Bozinovic, 2012). However, intraspecific variation in environmental tolerances and acclimation capacity have largely been overlooked in forecasts of species’ responses to climate change (Valladares et al., 2014; Bennett et al., 2019; Enriquez-Urzelai et al., 2020).

Elevational gradients constitute natural laboratories to investigate patterns of intraspecific variation in relation with environmental changes which occur over short spatial distances. Mountain habitats are characterised by steep environmental gradients over short geographic distances, such as a sharp temperature reduction with altitude (Körner, 2007) so that populations living at the upper and lower altitudinal extremes experience drastically different environmental conditions. Hence, species distributed along wide elevational gradients (i.e., from lowlands to the alpine belt of mountain systems) are likely to experience local adaptation or exhibit high levels of phenotypic plasticity (Keller et al., 2013). However, in elevation-generalist species with presumably high dispersal capacity, intraspecific variation in thermal traits will greatly depend on metapopulation dynamics, i.e., the degree of gene flow among populations, which might counteract trait divergence (e.g., Levy & Nufio, 2015). Indeed, the prevalence of adaptive clinal intraspecific variation in thermal tolerance with elevation remains unclear: whilst recent studies have revealed significant variation in thermal limits with elevation among populations of different taxa (e.g., Klok & Chown, 2003; Sørensen et al., 2005; Muñoz et al., 2014; Bishop et al., 2017; Senior et al., 2019; Enriquez-Urzelai et al. 2020) others have not found such elevational trend (Gvoždík & Castilla, 2001; Slatyer et al., 2016; Tonione et al., 2020).

In the case of freshwater species, most studies exploring thermal tolerance variation along environmental gradients have focused on interspecific comparisons and shown rather context-specific results depending on regions (temperate vs. tropical latitudes) and the specific taxa and tolerance trait analysed (e.g., Shah et al., 2017a,b; Pintanel et al., 2022). For example, using a comprehensive dataset of ectotherm species, Sunday et al. (2019) found that only lower thermal limits varied with elevation across freshwater species. In contrast, in a recent study, Carbonell et al. (under review) found that alpine species have both higher CTmax and lower CTmin than closely related species from lowlands. However, information on how populations within freshwater species vary in thermal tolerance with elevation is particularly scarce (but see Gutiérrez-Pesquera et al., 2022), and virtually non-existent for freshwater insects.

With regard to the factors driving variation in thermal tolerance across environmental gradients, two central hypotheses in thermal ecology are the mountain passes hypothesis (Janzen, 1967) and the climatic variability hypothesis (CVH; Stevens, 1989). Both hypotheses assume that organisms that have evolved in environments subject to higher thermal variability have greater thermal tolerance breadths and acclimation capacities than those from more stable environments. A corollary of these hypotheses is that organisms’ thermal limits are adapted to the climate extremes that they experience. The *Climate Extremes Hypothesis* (Pither, 2003) is a variant of the CVH, which predicts that extreme thermal events, even if rare, are a key selective agent in the evolution of thermal tolerance. Indeed, there is increasing evidence that rare but extreme thermal events play important roles in selecting for thermal tolerance (e.g., Smale & Wernberg, 2013; Kingsolver & Buckley, 2017; Coleman & Wernberg, 2020). Elevational gradients, varying strongly in daily and seasonal climatic regimes, provide an ideal arena for testing these hypotheses.

Here, we assess patterns of variation in critical thermal limits and their plasticity (acclimation capacity) in populations of a widespread aquatic beetle species (*Agabus bipustulatus* (Linnaeus, 1767), family Dytiscidae) across a 2,000-m elevational gradient, from lowland (ca. 400 masl) to alpine areas (ca. 2400 masl). This species represents an ideal study system to explore local adaption and plasticity of thermal traits in relation with the environmental challenges imposed by altitude, as it is a very common species widely distributed across the Western Palearctic region covering a large elevational range, from sea level to over 3000 masl. Indeed, this species shows a high intraspecific morphological variability across elevational gradients (Drotz et al., 2010), likely as a result of local adaption to different selective pressures. Accordingly, we also expect to find altitudinal clines in thermal tolerance. Specifically, as minimum, mean and maximum temperatures usually decline with elevation, and thermal physiology limits are driven by peak environmental temperatures (Overgaard et al., 2014; Buckley & Huey, 2016; Gutiérrez-Pesquera et al., 2016; Pintanel et al., 2019), we predict that higher elevation populations will show lower tolerances to high temperatures and higher tolerances to low temperatures, in agreement with the *Climate Extremes Hypothesis.* Based on previous studies on the plasticity of thermal limits in freshwater taxa (e.g. Shah et al., 2017a; Gutiérrez-Pesquera et al., 2022) we also explore whether acclimation capacity is positively related with climatic variability across populations at different elevations, in agreement with the CVH.

## MATERIAL AND METHODS

### Study system

*Agabus bipustulatus* is a predaceous, medium-large size diving beetle (9.0-11.0 mm), aquatic in all life stages, whose distribution covers the Western Palearctic region. We focused our study in five populations across the Iberian Peninsula (Figure 1, Table 1), where the species is widely distributed, mainly in lentic waters (Millán et al., 2014). The selected populations cover an elevation range from lowland (ca. 400 masl) to high-mountain areas (>2400 masl) in several Iberian mountain ranges, with decreasing mean, maximum and minimum temperatures along such gradient (Table 1).

**FIGURE 1.**
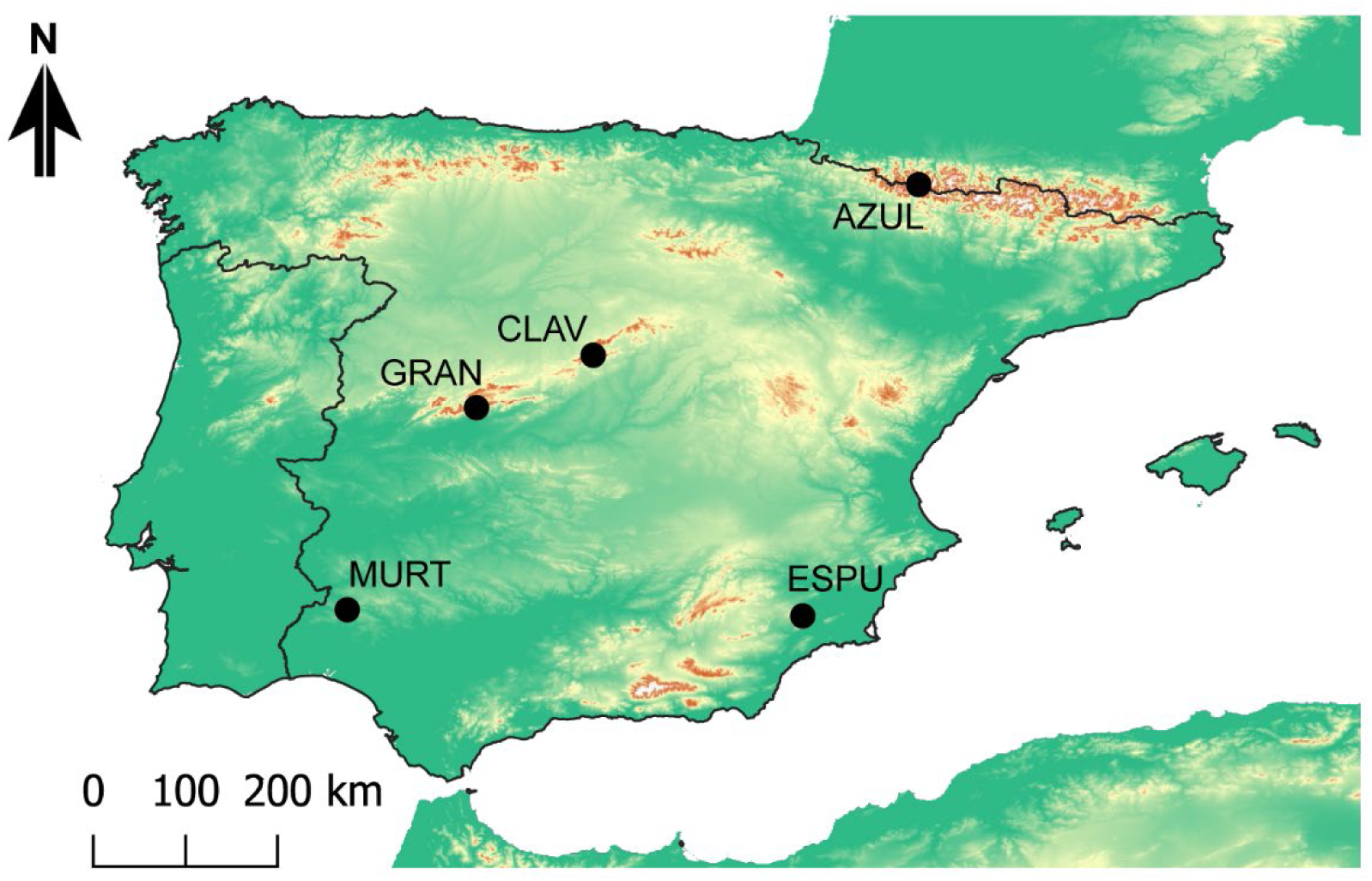
Study area and distribution of studied populations of *A. bipustulatus.* Population codes as in Table 1.

**TABLE 1.** Studied localities and climatic data (downloaded from WorldClim v. 2.1 database at 30 arc-s resolution, https://www.worldclim.org/)

| **Code** | **Name** | **Altitude**<br>**(masl)** | **Annual mean**<br>**temperature**<br>**(°C)** | **Min. annual**<br>**temperature**<br>**(°C)** | **Max. annual**<br>**temperature**<br>**(°C)** | **Mean diurnal**<br>**range**<br>**(°C)** | **Thermal**<br>**annual range**<br>**(°C)** |
| --- | --- | --- | --- | --- | --- | --- | --- |
| MURT | Río Múrtigas, La Nava, Huelva | 419 | 15.4 | 3.5 | 31.8 | 11.5 | 28.3 |
| ESPU | Fuente Blanca, Sierra Espuña | 1142 | 12.0 | -1.4 | 31.0 | 12.4 | 32.4 |
| GRAN | Laguna Grande de Gredos, Sierra de Gredos | 1943 | 6.1 | -4.3 | 21.9 | 7.7 | 26.2 |
| CLAV | Pools nr. Laguna de los Claveles, Sierra de Guadarrama | 2116 | 5.2 | -4.6 | 20.8 | 7.6 | 25.4 |
| AZUL | Ibón Azul Superior, Pyrenees | 2407 | 1.6 | -9.5 | 18.5 | 9.4 | 28.0 |

### Thermal tolerance experiments

Between 20-80 adult specimens per locality were collected alive in summer 2022 and transported within 24 h to the laboratory in 500 mL containers with moistened filter paper, placed in a portable refrigerator. Upon arrival, they were allowed to habituate to laboratory conditions for 3 days prior to experiments at 15 or 20 °C (i.e., temperatures approaching the temperature at collection in each site) and a 12:12 L:D photoperiod in a climatic chamber (SANYO MLR-351). Specimens were placed in 7 L aerated tanks with 2 L of bottled water from Sierra Nevada (Lanjarón ®), at densities lower than 50 individuals per litre and stones from the collection localities as substrate. Individuals were fed daily *ad libitum* with frozen chironomid larvae.

After the habituation period, groups of individuals were acclimated at 10, 15 or 20 °C for 7 days in climatic chambers, keeping the other conditions the same as before. In the population from *Río Múrtigas* (Table 1) only the 10°C acclimation treatment was performed due to insufficient sample size. Individuals were starved for 24 h prior to trials, as gut contents may modify thermal tolerance (Chown & Nicolson, 2004), and sets of individuals from each acclimation temperature were randomly divided in sub-groups of 10 beetles to estimate upper and lower thermal limits.

The heat tolerance of the studied populations was assessed by estimating the heat coma temperature (HCT) as the upper critical thermal limit (CTmax). HCT is defined as the temperature at which individuals experience paralysis prior to death, preceded by spasmodic movements of legs and antennae (Chown & Terblanche, 2006). It was estimated in air employing a dynamic method (Lutterschmidt & Hutchison, 1997) with a heating rate of 1 °C min^-1^, starting at the corresponding acclimation temperature. This is a standard ramping rate, widely used in thermal tolerance assays with arthropods (e.g., Diaz et al., 2002; Calosi et al., 2008; Wehner & Wehner, 2011). Specimens were gently dried with tissue paper and glued dorsally on a ceramic plate using adhesive tape to prevent escape during the trials. Trials were carried out in a controlled-temperature chamber (BINDER MK53. BINDER GmbH, Tuttlingen, Germany) coupled with an infrared thermographic camera (FLIR SC305, Teledyne FLIR, Wilsonville, Oregon, USA) to record body temperature. High quality images were also recorded with a video camera (Sony DCR-DVD110E, Sony Co., Tokyo, Japan) to determine accurately the moment of paralysis (cessation of movement of legs and antennae).

The cold tolerance of populations was estimated using the supercooling point (SCP) as a lower critical thermal limit (CTmin). SCP is the temperature at which the body fluids of the organism begin to freeze when specimens are exposed to cooling. SCP was estimated as the lower temperature reached before the release of the latent heat of crystallization, employing also a dynamic method with a cooling rate of -1 °C min^-1^ and infrared thermography. All specimens were sexed after experiments. Thermal images were analysed with the software ThermaCAM Researcher Professional v. 2.10 (FLIR Advanced Thermal Solutions; ATS; Croissy-Beaubourg, France).

For each population and acclimation treatment, we estimated the thermal breadth as the difference between average HCT and SCP. Plasticity of upper and lower thermal limits was calculated as the acclimation response ratio (ARR), which describes the change in thermal limits with a given change in acclimation temperature (Gunderson & Stillman, 2015). We used values of HCT and SCP averaged per acclimation treatment and estimated ARRs as ARR = Δ HCT or Δ SCP /Δ°C.

### Data analysis

In order to assess whether HCT and SCP varied among populations and acclimation treatments and whether acclimation effects differ among populations, we used generalised linear models (GLMs) with a normal error structure and identity link function, with temperature (10, 15 and 20°C) and population (and their interaction) as fixed factors. Sex was initially included as a fixed factor and removed if not significant. For significant predictors, we performed posthoc pairwise comparisons with Bonferroni adjustment for P-values to identify significant differences between specific pairs of populations, temperatures or population x temperature combinations.

We assessed the relationships between thermal tolerance limits (HCT and SCP) and environmental predictors (elevation, average, minimum and maximum annual temperature and diurnal and annual thermal range) using simple linear regressions. Climatic data were obtained from WorldClim v. 2.1 (Fick & Hijmans, 2017) layers at 30 arc-s resolution (ca. 1km at the Equator). We first checked for correlation between predictors and selected a subset of uncorrelated variables (Pearson coefficient < 0.85) to perform the regressions.

Finally, since the results obtained prevented us for testing the relationship between plasticity of the thermal limits (ARR values) and thermal variability (see results), we only compared ARR values among populations in a descriptive way.

All analyses were performed in R v. 4.1.2 (R Core Team, 2021).

## RESULTS

### Effects of population and acclimation treatments on critical thermal limits

Heat coma temperatures differed significantly among populations (Table 2). Posthoc tests revealed significant differences only between CLAV and ESPU populations (χ2 = 12.882 P_1,127_ = 0.003; Figure 2, Supplementary Material: Table S1) and such differences varied between acclimation treatments (population x acclimation interaction; Table 2), being significant only in the 15°C treatment (χ2 = 17.933, P_1,110_ < 0.001; Figure 2, Supplementary Material: Table S1). Females showed higher HCTs than males (Table 2).

**TABLE 2.**
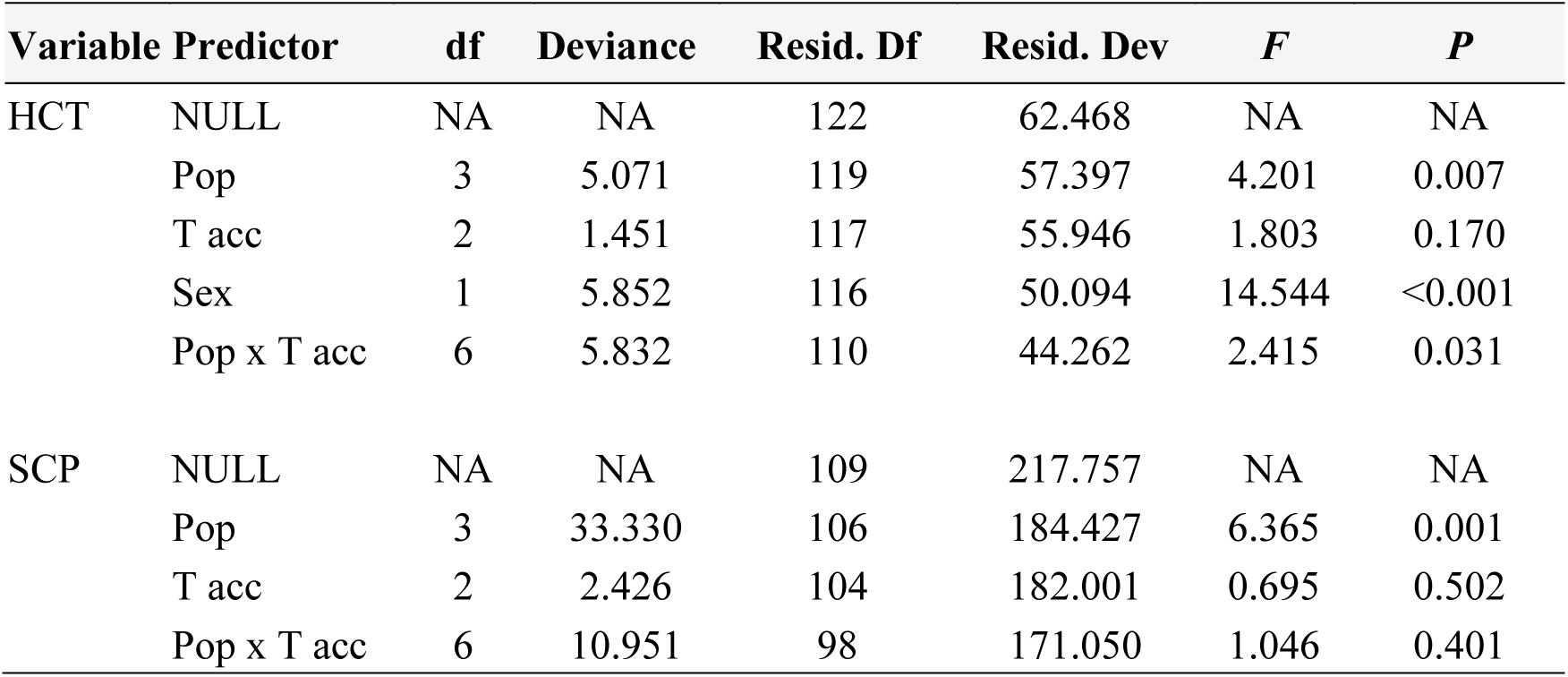
Results of GLMs assessing the effects of population (Pop), acclimation temperature (T acc) and sex on heat coma temperatures (HCT) and supercooling points (SCP).

**FIGURE 2.**
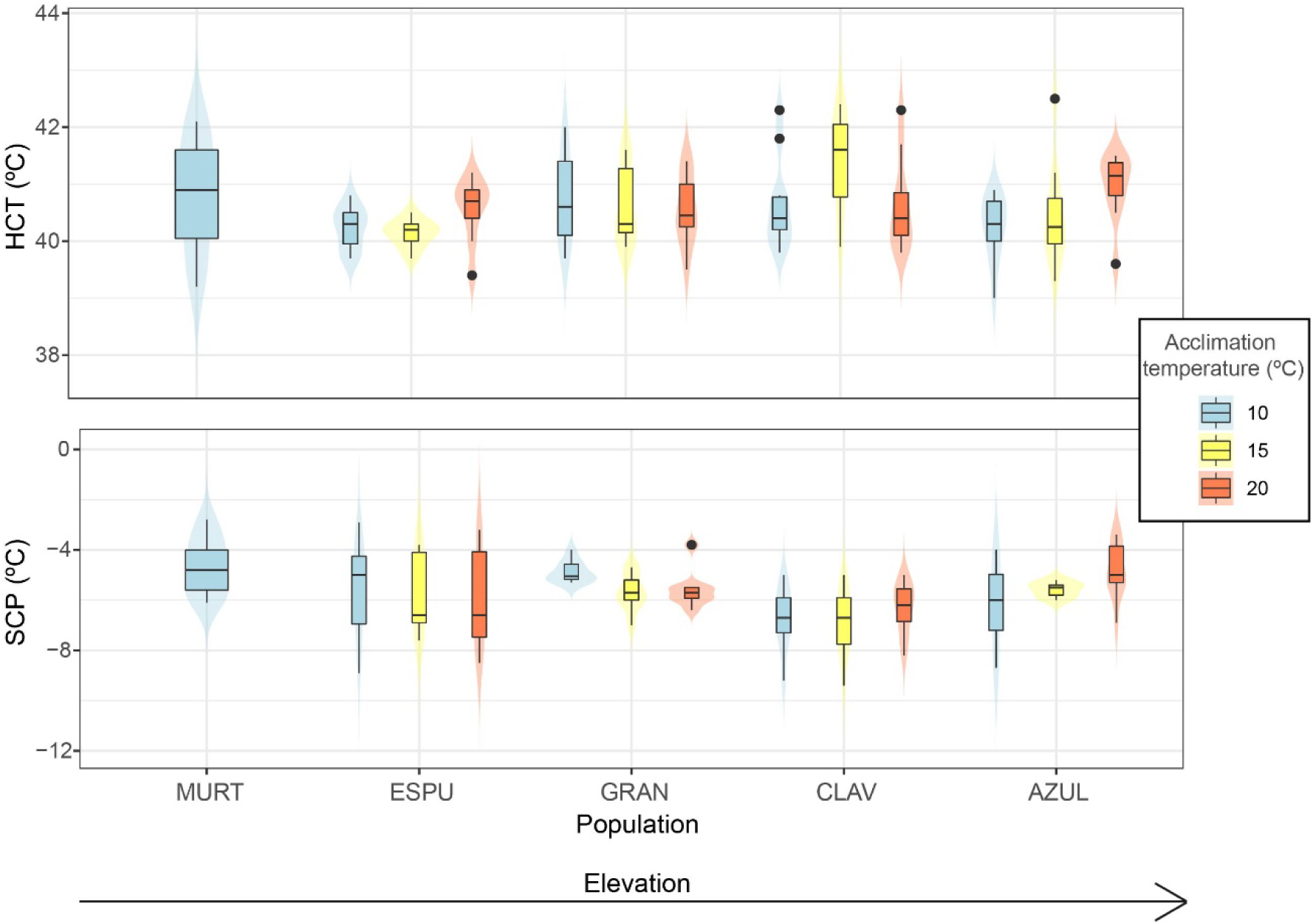
Violin plots with boxplot overlay with heat coma temperature (HCT, upper panel) and supercooling point (SCP, lower panel) for each population and acclimation temperature. Boxplots represent Q25, median and Q75, whiskers are Q10 and Q90 and dots are outliers. The violin plot lines indicate a kernel density of the distribution (i.e., a smoothed histogram). Population codes as in Table 1.

We found significant differences in supercooling points between populations, but not significant effects or acclimation temperature or sex (Table 2). The CLAV population showed significantly lower SCPs than the remaining populations (all Ps < 0.05 in posthoc tests; Figure 2, Supplementary Material: Table S2).

### Relationships between thermal tolerance limits and environmental predictors

After checking for correlations among environmental variables (Figure S1), we selected altitude and annual thermal range as predictors for simple linear regressions. We found no significant relationships between any thermal tolerance trait and environmental predictors, except for a weak, yet significant negative correlation between HCT and annual thermal range (Table 3, Figure 3).

**TABLE 3.** Results of the linear regressions between heat coma temperature (HCT) or supercooling point (SCP) and environmental variables.

| Dependent variable | Predictor | Coefficient±SE | t value | P | R<sup>2</sup> |
| --- | --- | --- | --- | --- | --- |
| HCT | Altitude | Intercept: 40.45±0.20 | 200.686 | <0.001 | -0.003 |
|  |  | Altitude: 0.00008±0.0001 | 0.799 | 0.426 |  |
|  | Annual thermal range | Intercept: 42.61±0.67 | 63.435 | <0.001 | 0.055 |
|  |  | Annual thermal range: -0.072±0.024 | -2.994 | **0.003** |  |
| SCP | Altitude | Intercept: -5.073±0.402 | -12.623 | <0.001 | 0.016 |
|  |  | Altitude: -0.004±0.0002 | -1.696 | 0.092 |  |
|  | Annual thermal range | Intercept: -7.552±1.325 | -5.701 | <0.001 | 0.008 |
|  |  | Annual thermal range: 0.065±0.047 | 1.391 | 0.167 |  |

**FIGURE 3.**
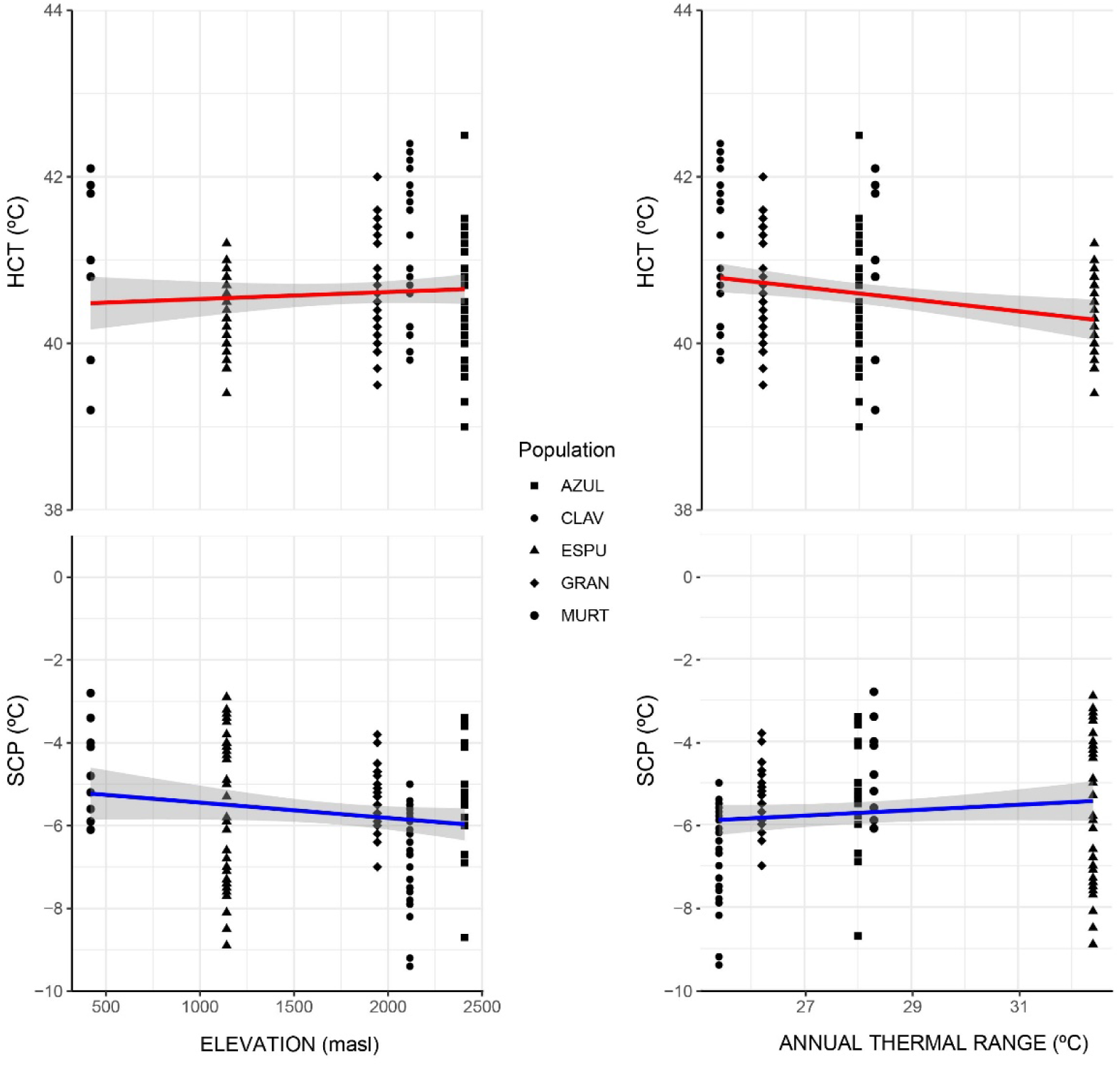
Simple linear regressions between heat coma temperature (HCT) and supercooling point (SCP) and environmental variables. Regression lines and 95% confidence intervals are shown. Population codes as in Table 1.

### Thermal breadth and Acclimation Response Ratios

Thermal breadths ranged between 45.48 °C (MURT population, at the lowest altitude) and 47.55 °C (CLAV population, at 2216 masl) (Table 4). ARR values for HCTs and SCPs were low in general (<0.1; Table 4). Populations with lower daily and annual thermal variability (Table 1) show a slightly higher thermal plasticity (i.e., CLAV and AZUL showed ARR values of the HCT > 0.1 for some acclimation temperature pairs and the same happened with GRAN and AZUL for SCP; Table 4)

**TABLE 4.** Thermal breadths and acclimation response ratios (ARR) for heat coma temperatures (HCT) and supercooling points (SCP) for each population. Population codes as in Table 1.

|  | **MURT** | **ESPU** | **GRAN** | **CLAV** | **AZUL** |
| --- | --- | --- | --- | --- | --- |
| Thermal breadth (°C) | 45.48 | 46.06 | 46.00 | 47.55 | 46.06 |
| HCT ARR 10-15°C |  | 0.018 | 0.019 | 0.154 | 0.037 |
| ARR 15-20°C |  | 0.087 | 0.010 | 0.158 | 0.111 |
| ARR 10-20°C |  | 0.034 | 0.014 | 0.002 | 0.074 |
| SCP ARR 10-15°C |  | 0.040 | 0.160 | 0.031 | 0.119 |
| ARR 15-20°C |  | 0.051 | 0.068 | 0.044 | 0.138 |
| ARR 10-20°C |  | 0.040 | 0.160 | 0.031 | 0.119 |

## DISCUSION

Intraspecific variation in physiological traits is common in broadly distributed species (e.g., Lardies et al., 2010; Barria & Bacigalupe, 2017; Maebe et al., 2021). Populations exposed to contrasting climate extremes and thermal variability along elevational gradients are prone to be subject to selection on thermal tolerance traits and physiological plasticity, in agreement with the CVH and the *Climate Extremes Hypothesis* (Keller et al., 2013). Here, we found significant variation in thermal limits among populations of the widespread diving beetle *A. bipustulatus*, but no evidence that such intraspecific variation could be related to local adaptation to different thermal conditions along altitudinal gradients. In contrast, the studied populations did not differ in the plasticity of thermal limits, which was overall and consistently low in all populations.

The significant variation in upper (HCT) and lower (SCP) thermal limits among *A. bipustulatus* populations was mainly attributed to the higher heat and cold tolerance displayed by one specific population, located at high altitude (CLAV population, in *Sierra de Guadarrama*, at 2116 masl), with respect to the other ones. Thermal limits were not correlated to elevation or climatic variables, except from a significant but weak negative correlation between HCT and the annual thermal range. We cannot provide at present an ecological explanation for such counter-intuitive relationship, which contradicts the CVH. Furthermore, it cannot be dismissed that the relatively coarse air temperature variables used here as climatic predictors may not accurately reflect the actual thermal variations experienced by the populations (see below). However, it is apparent that our data did not support the CHV or the *Climatic Extremes Hypotheses*. The lack of a clear clinal variation in thermal tolerance limits in *A. bipustulatus* suggests conservatism of the thermal niche and is consistent with the findings of Gutiérrez-Pesquera et al. (2022), which also found no relationship between thermal limits and elevation or temperature across populations of *Rana parvipalmata* tadpoles. In contrast, other studies have reported significant relationships between thermal tolerance traits and altitude within species of different terrestrial and semiaquatic taxa (e.g., salamanders and lizards — Trochet et al., 2018, frogs —Enriquez-Urzelai et al. 2020 or ants —Tonione et al. 2020). Future studies of intraspecific variation of thermal tolerance along environmental gradients in other freshwater invertebrates would help us understand to which extent the thermal niche conservatism observed here is common among freshwater species.

Thermal acclimation may prevent directional selection on other physiological thermal traits, constraining the adjustment of thermal tolerance ranges to the local environment (Chevin et al., 2010; Huey et al., 2012). According to the CVH, we expected that populations living in more variable thermal environments would exhibit greater plasticity in their thermal tolerances. However, any of the studied populations of *A. bipustulatus* showed significant acclimation capacity of either HCP or SCP along the acclimation temperatures tested, and AAR values were generally low. Although plastic thermal tolerances may be adaptive (Sultan & Spencer, 2002), it has been widely discussed that plasticity has costs and evolutionary constraints (e.g., Dewitt et al., 1998; Relyea, 2002; Auld et al., 2010; Murren et al., 2015). The apparent lack of thermal acclimation capacity in *A. bipustulatus* is consistent with previous studies, in which several dytiscid beetles, including this species and other congeners, have been shown to have in general limited or no acclimation capacity in several thermal tolerance traits (Pallarés et al., 2020; Pallarés et al., 2024; Carbonell et al., under review). In contrast, plasticity of upper and lower thermal limits has been shown to vary notably between species of other lineages of this family, as the genus *Deronectes* (Calosi et al., 2010). These contrasting results stress the need to improve our understanding of the factors shaping thermal tolerance traits in freshwater insects.

The relative low variation in thermal limits and their plasticity between *A. bipustulatus* populations might be explained by different mechanisms. First, in a widespread species with a pressumably high dispersal capacity such as *A. bipustulatus* (Lundkvist et al., 2002; Drotz et al., 2010; Bilton, 2023), gene flow may erase to some extent divergence of thermal physiology between populations, according with the hypothesis of ‘ephemeral divergence’ (Futuyma, 2010). Environmental transitions along altitudinal gradients typically occur at spatial scales that are small relative to the dispersal distances of many species. Then, the effects of divergent selection may be opposed by gene flow, which, if strong enough, acts to homogenize allele frequencies between environments (Lenormand, 2002; Keller et al., 2013; Bachmann et al., 2020). However, an increasing number of studies have provided evidence of maintenance of local adaptation despite gene flow (Tigano & Friesen, 2016).

Alternatively, environmentally driven adaptations in other traits may buffer the degree of climatic variation and exposure to climate extremes, lowering selective pressures for population differentiation in physiological thermal tolerance. For example, differences in reproductive phenology between populations may reduce the actual differences in temperatures experienced by adults across different elevations. It has been recently suggested that the endemic alpine diving beetle *A. nevadensis*, closely related to *A. bipustulatus*, might overwinter in alpine lakes in the larval stage (Carbonell et al., 2024). This might be also the case of high-altitude populations of *A. bipustulatus*, so that adults might not been actually exposed to extreme cold temperatures and selection on adult cold tolerance could be relaxed. Also, microhabitat selection might allow adults to avoid peak winter and summer temperatures, to some extent. The potential for behavioural plasticity could limit the evolution of plasticity in physiological traits, which is known as the Borgert effect (Bogert, 1949; Huey et al., 2003). Although the scope for behavioural thermoregulation is limited in the aquatic environment, adults of this species (whose bimodal respiration allows them to breadth from atmospheric oxygen), could use wet shelters in river and lake banks or in land under rocks, mud or vegetation, buffered from extreme air temperatures. Adults of alpine populations of related dytiscids have been observed under moss in pond shores in high densities (P. Abellán & J.A. Carbonell, pers. obs.). In relation to this point, it should be noted that most animals experience climate at fine-scale patches and indeed, increasing evidence suggests that macroclimatic variables may only weakly predict tolerance limits and physiological niches compared with microclimatic ones (e.g. Pintanel et al., 2019, Farallo et al., 2020; Bartolini & Giomi, 2021). Thus, exploring the relationship between thermal tolerance traits and relevant microclimatic variables in aquatic insects is another important avenue of future research.

In summary, our results suggest that *A. bipustulatus* has a limited adaptive potential and plasticity of thermal tolerance, despite being a species distributed along a wide environmental gradient, including extreme alpine habitats, where the environment imposes substantial constraints. These results highlight the need to improve our current understanding of how populations of aquatic insects vary in thermal tolerance with elevation, and explore the role of other mechanisms (e.g., phenological adaptations, behavioural thermoregulation) in coping with climatic constraints. In a context of climate change, a low capacity to adjust thermal tolerance to local and changing environmental conditions trough adaptive changes and physiological plasticity could imply a limited resilience to global warming of the study species. Previous work on related species has shown that their heat tolerance limits are above the predicted maximum temperatures in their localities (Pallarés et al., 2020). However, that study also stressed that the capacity to adjust such limits through acclimation is limited, as we found here in *A. bipustulatus*. Indeed, our results are in line with previous work suggesting that the thermal plasticity of ectotherms might have limited potential to buffer them from global warming (Gunderson & Stillman, 2015). In such a context, identifying those populations which will be more exposed to warming from climate change should be a priority.

## ACKNOWLEDGEMENTS

This work was supported by the R&D project id. PID2019-108895GB-I00, funded by MCIN/AEI/ 10.13039/501100011033 and VI PPIT Universidad de Sevilla (IV.7 Ayuda Suplementaria a Grupos de Investigación por captación de fondos en las convocatorias de proyectos de investigación del Plan Estatal; 2020/1110). SP was funded by a postdoctoral contract from the ‘Consejería de Economía, Conocimiento, Empresas y Universidad de la Junta de Andalucía-Fondo Social Europeo de Andalucía 2014-2020’ (‘Talento Doctores, PID 2020’ program, grant id. SP-DOC_01211). JAC was funded by a postdoctoral contract from the ‘María Zambrano’ program (grant id. 19868), by the Spanish ‘Ministerio de Universidades’ (funded by European Union - NextGenerationEU). FP was funded by a postdoctoral contract from the ‘Consejería de Economía, Conocimiento, Empresas y Universidad de la Junta de Andalucía-Fondo Social Europeo de Andalucía 2014-2020’ We are especially grateful to David Sánchez-Fernández for his help at various stages of this project.

## Notes

### Competing Interest Statement

The authors have declared no competing interest.

