## Supplementary Material for "Intraspecific variation of thermal tolerance in freshwater insects along elevational gradients: the case of a widespread diving beetle"

**SUPLEMENTARY MATERIAL**

**TABLE S1**. Posthocs pairsiwe comparisons of heat coma temperature between pairs of populations (Pop) and population x acclimation temperature (T acc) combinations. Population codes as in Table 1.

| Factor | Compared pairs | Value | df | $\chi^2$ | *P* |
|---|---|---|---|---|---|
| **Pop** | AZUL-CLAV | -0.314 | 1 | 3.344 | 0.674 |
| | AZUL-ESPU | 0.345 | 1 | 3.613 | 0.573 |
| | AZUL-GRAN | -0.004 | 1 | 0.001 | 1.000 |
| | AZUL-MURT | -0.156 | 1 | 0.393 | 1.000 |
| | CLAV-ESPU | 0.659 | 1 | 12.882 | 0.003 |
| | CLAV-GRAN | 0.31 | 1 | 3.239 | 0.719 |
| | CLAV-MURT | 0.158 | 1 | 0.402 | 1.000 |
| | ESPU-GRAN | -0.349 | 1 | 3.807 | 0.511 |
| | ESPU-MURT | -0.501 | 1 | 3.898 | 0.483 |
| | GRAN-MURT | -0.152 | 1 | 0.378 | 1.000 |
| | Residuals | NA | 127 | NA | NA |
| **T acc x Pop** | 10-15: AZUL | -0.183 | 1 | 0.501 | 1.000 |
| | 10-20: AZUL | -0.674 | 1 | 6.115 | 0.161 |
| | 15-20: AZUL | -0.490 | 1 | 3.239 | 0.863 |
| | 10-15: CLAV | -0.618 | 1 | 4.618 | 0.380 |
| | 10-20: CLAV | 0.020 | 1 | 0.005 | 1.000 |
| | 15-20: CLAV | 0.638 | 1 | 4.921 | 0.318 |
| | 10-15: ESPU | 0.092 | 1 | 0.107 | 1.000 |
| | 10-20: ESPU | -0.584 | 1 | 2.989 | 1.000 |
| | 15-20: ESPU | -0.676 | 1 | 3.644 | 0.675 |
| | 10-15: GRAN | 0.105 | 1 | 0.141 | 1.000 |
| | 10-20: GRAN | 0.187 | 1 | 0.444 | 1.000 |
| | 15-20: GRAN | 0.082 | 1 | 0.099 | 1.000 |
| | Residuals | NA | 110 | NA | NA |
| **Pop x Tacc** | AZUL-CLAV: 10 | -0.391 | 1 | 2.075 | 1.000 |
| | AZUL-ESPU: 10 | 0.135 | 1 | 0.264 | 1.000 |
| | AZUL-GRAN: 10 | -0.394 | 1 | 1.961 | 1.000 |
| | CLAV-ESPU: 10 | 0.526 | 1 | 3.699 | 0.980 |
| | CLAV-GRAN: 10 | -0.002 | 1 | 0.000 | 1.000 |
| | ESPU-GRAN: 10 | -0.528 | 1 | 3.558 | 1.000 |
| | AZUL-CLAV: 15 | -0.826 | 1 | 8.876 | 0.052 |
| | AZUL-ESPU: 15 | 0.41 | 1 | 2.101 | 1.000 |
| | AZUL-GRAN: 15 | -0.105 | 1 | 0.163 | 1.000 |
| | CLAV-ESPU: 15 | 1.236 | 1 | 17.933 | <0.001 |
| | CLAV-GRAN: 15 | 0.721 | 1 | 6.986 | 0.148 |
| | ESPU-GRAN: 15 | -0.515 | 1 | 3.384 | 1.000 |
| | AZUL-CLAV: 20 | 0.302 | 1 | 1.132 | 1.000 |
| | AZUL-ESPU: 20 | 0.224 | 1 | 0.410 | 1.000 |
| | AZUL-GRAN: 20 | 0.466 | 1 | 2.934 | 1.000 |

| | | | | |
|---|---|---|---|---|
| CLAV-ESPU: 20 | -0.078 | 1 | 0.050 | 1.000 |
| CLAV-GRAN: 20 | 0.164 | 1 | 0.361 | 1.000 |
| ESPU-GRAN: 20 | 0.243 | 1 | 0.516 | 1.000 |
| Residuals | NA | 110 | NA | NA |

**TABLE S2**. Posthocs pairsiwe comparisons of supercooling points between pairs of populations (codes as in Table 1).

| **Compared pairs** | **Value** | **df** | $\chi^2$ | ***P*** |
|---|---|---|---|---|
| AZUL-CLAV | 1.372 | 1 | 13.375 | 0.003 |
| AZUL-ESPU | 0.446 | 1 | 1.429 | 1.000 |
| AZUL-GRAN | 0.078 | 1 | 0.041 | 1.000 |
| AZUL-MURT | -0.614 | 1 | 1.370 | 1.000 |
| CLAV-ESPU | -0.926 | 1 | 7.897 | 0.050 |
| CLAV-GRAN | -1.294 | 1 | 14.120 | 0.002 |
| CLAV-MURT | -1.986 | 1 | 16.087 | 0.001 |
| ESPU-GRAN | -0.367 | 1 | 1.156 | 1.000 |
| ESPU-MURT | -1.06 | 1 | 4.614 | 0.317 |
| GRAN-MURT | -0.693 | 1 | 1.893 | 1.000 |
| Residuals | NA | 114 | NA | NA |

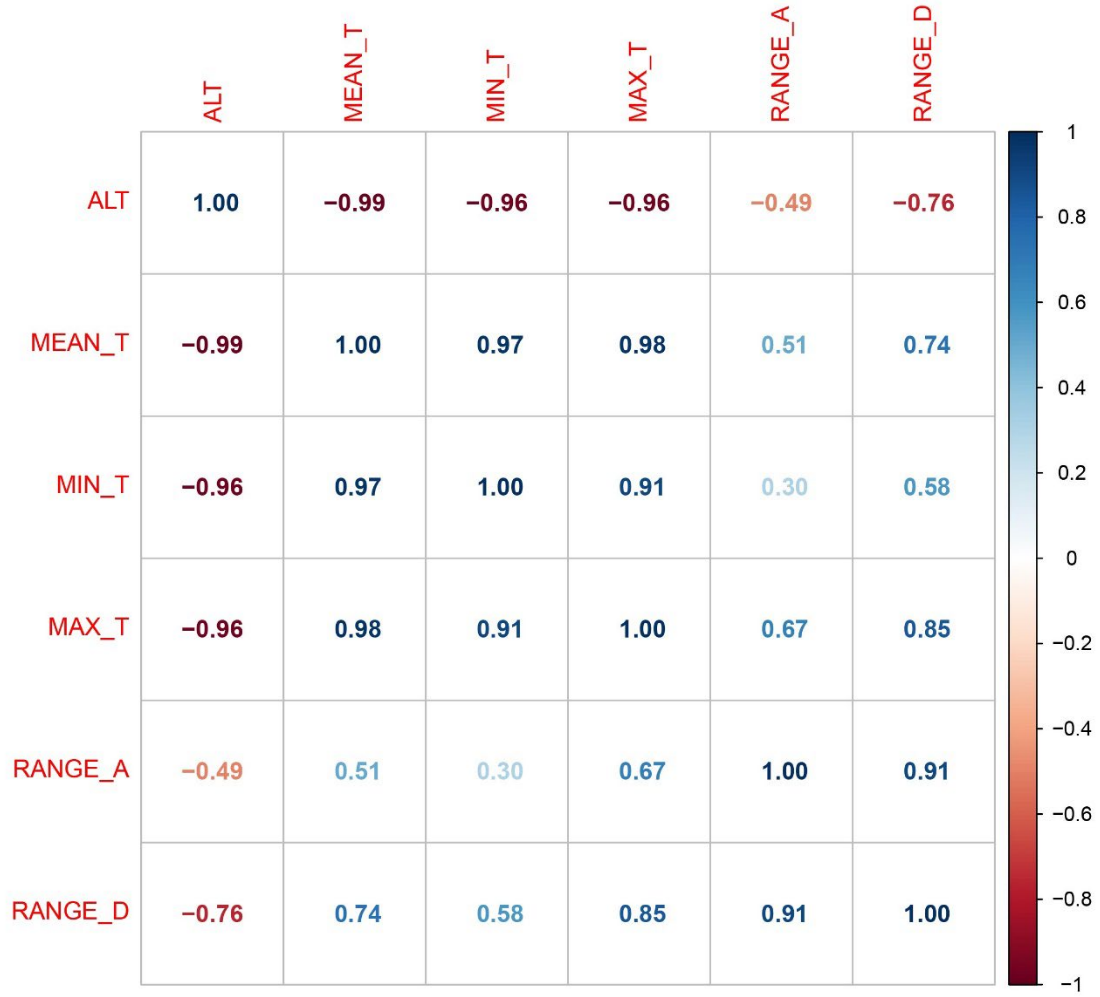

**FIGURE S1**. Correlogram showing Pearson coefficients among environmental variables: altitude (ALT), annual mean (MEAN_T), minimum (MIN_T) and maximum (MAX_T) temperatures, annual (RANGE_A) and daily (RANGE_D) thermal ranges.
